# Stop Bickering! Reconciling Signaling Pathway Databases with Network Topologies

**DOI:** 10.1101/2021.08.03.454954

**Authors:** Tobias Rubel, Pramesh Singh, Anna Ritz

**Affiliations:** Biology Department, Reed College, Portland, Oregon, USA

**Keywords:** Signaling Pathways, Biological Networks, Network Topology, Graphlets

## Abstract

A major goal of molecular systems biology is to understand the coordinated function of genes or proteins in response to cellular signals and to understand these dynamics in the context of disease. Signaling pathway databases such as KEGG, NetPath, NCI-PID, and Panther describe the molecular interactions involved in different cellular responses. While the same pathway may be present in different databases, prior work has shown that the particular proteins and interactions differ across database annotations. However, to our knowledge no one has attempted to quantify their structural differences. It is important to characterize artifacts or other biases within pathway databases, which can provide a more informed interpretation for downstream analyses. In this work, we consider signaling pathways as graphs and we use topological measures to study their structure. We find that topological characterization using graphlets (small, connected subgraphs) distinguishes signaling pathways from appropriate null models of interaction networks. Next, we quantify topological similarity across pathway databases. Our analysis reveals that the pathways harbor database-specific characteristics implying that even though these databases describe the same pathways, they tend to be systematically different from one another. We show that pathway-specific topology can be uncovered after accounting for database-specific structure. This work present the first step towards elucidating common pathway structure beyond their specific database annotations.

## 1. Introduction

Cells respond to signals through a series of molecular interactions, culminating in gene expression changes that alter the cell’s behavior. The protein-protein interactions that occur in response to specific stimuli are described as signaling pathways. These pathways characterize cell growth, proliferation, stress, death, and transport, among many other biological processes. For over a decade, the growing knowledge about cellular signaling has been collected in databases such as KEGG,^13^ NetPath,^12^ Reactome,^9^ NCI-PID,^22^ and Panther.^15^ These resources are all manually curated, organized into specific signaling pathways, and comprise protein interactions supported by scientific literature. They are the scientific community’s best guess as to how proteins interact within a larger system of cellular response and are often the starting point for many downstream analyses of ‘omic data such as gene function enrichment and identifying genetic associations. Pathway databases have also seen extensive use for studying human diseases – many databases focus on pathways that are known to be dysregulated by disease^2,12,22^ or describe the altered pathways themselves.^9,13,26^

While the number and utility of signaling pathway databases grows, there still remain limitations in their broad use. Signaling pathways from different databases are often incomplete,^3,5,19^ though they contain high-quality interactions due to the database’s manual curation steps. Pathways with the same name may contain different protein-protein interactions or pathways characterizing the same response may be called different names.^3,7,24^ These challenges arise for a number of reasons: pathway nomenclature has not been standardized, pathway crosstalk and noncanonical signaling blurs the pathway boundaries, and we simply have not yet quantified all of the biological interactions that occur. The lack of consistency across pathway databases indicates that the choice of database can change the results of down-stream ‘omic analysis, which has been previously shown.^16^ New databases integrate existing pathways and offer standardized APIs and data file formats.^16,17,20,23,25^

However, we are still left with a fundamental question: *How do we reconcile signaling pathway annotations across databases*? Reconciling pathways is different from integrating pathways, which has been the focus of related endeavors. Work on protein interaction networks have shown that simply taking the union of the networks is prone to propagating noise.^11^ Instead, we consider the databases separately and strive to elucidate pathway-specific features that are shared across databases. Our working hypothesis is that, even though each database is manually curated with different goals and scopes, if they are describing similar signaling pathways then we should be able to uncover some information about the pathway structure.

We represent signaling pathways as graphs, allowing us to leverage the considerable theory developed for characterizing networks. There are lots of ways to characterize the structure of networks, ranging from extremely simple (and interpretable) summary statistics like degree distribution to more expressive measures. Signaling pathways have been analyzed using topological features such as degree, clustering coefficient, and centralities.^29,30^ However, despite their virtues, these statistics are too simple to fully characterize complex networks. Topological structures called graphlets^18,28^ have been shown to characterize networks better than simple summary statistics, and are still easy to interpret. Graphlets are small, connected subgraphs that have been used to analyze empirical networks such as world trade networks, social networks, and protein interaction networks. Two- and three-node graphlets have also been used to derive global and local network statistics that are robust to network size.^8^ Graphlet-based measures are useful characterizations of networks beyond node degree, clustering coefficient, or other centralities.

### Contributions

This paper is organized in three parts. First, we describe our methodology for collecting, parsing, and representing signaling pathways. Using graphlet-based network embeddings, we then examine pathway topologies in databases compared to suitable controls, and find that pathways are distinguishable from null models. Finally, we compare pathway databases to one another. For this last part, we identify similar pathways across databases which we call *corresponding pathways*. However, we find that pathways cluster by databases rather than by corresponding pathways which indicates that databases contain consistent topological structure and potentially obfuscates shared structure among pathway annotations. Using a regression framework, we correct the database structure and reveal pathway-specific topological structure where corresponding pathways cluster together. These results collectively indicate that, while pathway databases are manually curated with different scopes and intentions, the same pathway shares topological features across databases.

## 2. Data Collection and Processing

We considered ten pathway databases for this analysis, which all contain manually-curated human pathways grouped by phenotype or response. Our goal was to select pathway databases that were similar orders of magnitude in size, had a broad focus on different types of signaling, and did not contain other databases as subsets. We chose seven pathway databases from this list (Table 1). We excluded Reactome^9^ after finding that the parsed pathways were much larger than the others (Supplementary Fig. S1), we excluded CausalBioNet^2^ due to its focus on pulmonary and vascular signaling, and we excluded WikiPathways^23^ since it combines multiple pathway databases.

**Table 1.** Pathway databases used in this analysis, after converting pathways to undirected graphs and removing pathways with fewer than ten interactions. Number of pathways, mean and standard devation shown. The *\*-expanded* datasets convert complexes and families into protein identifiers (see Section 2.1).

| Database | Focus | Parse Source | $n$ | # Nodes | # Edges |
| --- | --- | --- | --- | --- | --- |
| INOH<sup>27</sup> | Hierarchical Model | PathwayCommons<sup>20</sup> | 114 | $30 \pm 47.00$ | $313 \pm 1063.06$ |
| KEGG<sup>13</sup> | Broad Focus | KEGG<sup>13</sup> | 243 | $38 \pm 30.67$ | $40 \pm 30.80$ |
| *KEGG-expanded*<sup>13</sup> | Broad Focus | KEGG<sup>13</sup> | 268 | $70 \pm 59.50$ | $252 \pm 417.68$ |
| NetPath<sup>12</sup> | Immune & Cancer | NetPath<sup>12</sup> | 32 | $73 \pm 61.67$ | $134 \pm 159.68$ |
| Panther<sup>15</sup> | Primary Signaling | PathwayCommons<sup>20</sup> | 93 | $57 \pm 50.02$ | $389 \pm 620.55$ |
| PathBank<sup>26</sup> | Model Organisms | Pathbank<sup>26</sup> | 576 | $21 \pm 29.18$ | $148 \pm 543.32$ |
| PID<sup>22</sup> | Cancer | PathwayCommons<sup>20</sup> | 203 | $102 \pm 78.22$ | $510 \pm 788.72$ |
| SIGNOR<sup>14</sup> | Binary Causal | NDEx<sup>17</sup> | 36 | $25 \pm 7.76$ | $39 \pm 31.08$ |
| *SIGNOR-expanded*<sup>14</sup> | Binary Causal | NDEx<sup>17</sup> | 36 | $38 \pm 24.20$ | $169 \pm 483.34$ |
| Interactome | Broad Focus | PathwayCommons<sup>20</sup> | 1 | 18494 | 1017054 |

### 2.1. From Pathways to Undirected Graphs

We strove to parse the databases from pathway compendia such as PathwayCommons^20^ and NDEx,^17^ which offer APIs and a unified file format. However, some pathways were parsed from the original source. While many of these databases are actively maintained, some resources such as NCI-PID and NetPath are no longer updated yet still contain useful information. We also parsed all of Pathway Commons, which includes experimentally-sourced interaction databases, as the interactome that is used to generate null models in Section 3.2 (Table 1).

To topologically characterize signaling pathways, we intentionally started simple. We converted every pathway into an undirected graph by parsing Simple Interaction Format (SIF) files. These files were pulled directly from PathwayCommons or were converted from BioPAX format using PaxTools. We only considered interactions that involved proteins and required pathways to contain at least ten undirected edges. Many metabolic networks, for example, were ignored due to this requirement. We mapped all proteins into HGNC namespace using the HGNC mapper (https://www.genenames.org/download/custom/).

Two databases, KEGG and SIGNOR, capture protein families and protein complexes in their networks.^13,14^ For these databases, we parsed a collapsed version which includes complexes and families as nodes in the network and an expanded version that converts such entities into their constitutive proteins. Interactions that include families or proteins were expanded to add an edge for every protein member (e.g., a two-protein family connected to a three-protein complex added six undirected edges). Further, protein complexes were connected in an “all-vs-all” manner to indicate physical interaction (e.g., a three-protein complex added three undirected edges). As expected, the average number of nodes and edges is larger for the expanded versions of the KEGG and SIGNOR databases (Table 1 and Supplementary Fig. S1). In total, we considered 1, 592 pathways in nine datasets that captured pathways from seven distinct databases.

### 2.2. Topological Characterization of Networks

Graphs have been characterized by global and local characteristics such as degree distribution, clustering coefficient, and path-based centralities. Topological structures such as network motifs (small connected subgraphs) have been shown to characterize networks. A natural generalization of a network motif is to enumerate all possible connected graphs of a specific size. This collection of networks have been coined as *graphlets*.

#### Graphlets

Graphlets, first introduced by Przulj et al.,^18^ are an enumeration of small, connected non-isomorphic graphs. We focus on undirected graphlets with up to five nodes (Fig. 1A). Graphlets can be efficiently computed, as described in other work.^1,10^ Here we provide an intuitive description of this calculation. An *automorphism orbit* (or *orbit* for short) of a graphlet is a set of nodes that are automorphic within a specific graphlet. A graphlet’s orbits summarize the possible distinct positions of each node in each graphlet. For example, there is one orbit in *G*_0_, two orbits in *G*_1_ (the middle node and the outer nodes), and one orbit in *G*_3_. There are a total of 73 orbits in the 30 graphlets in Fig. 1A). Once these orbits have been counted, they can be combined to produce graphlet counts for the network. We use ORCA to count orbits for each node.^10^

**Fig. 1.**
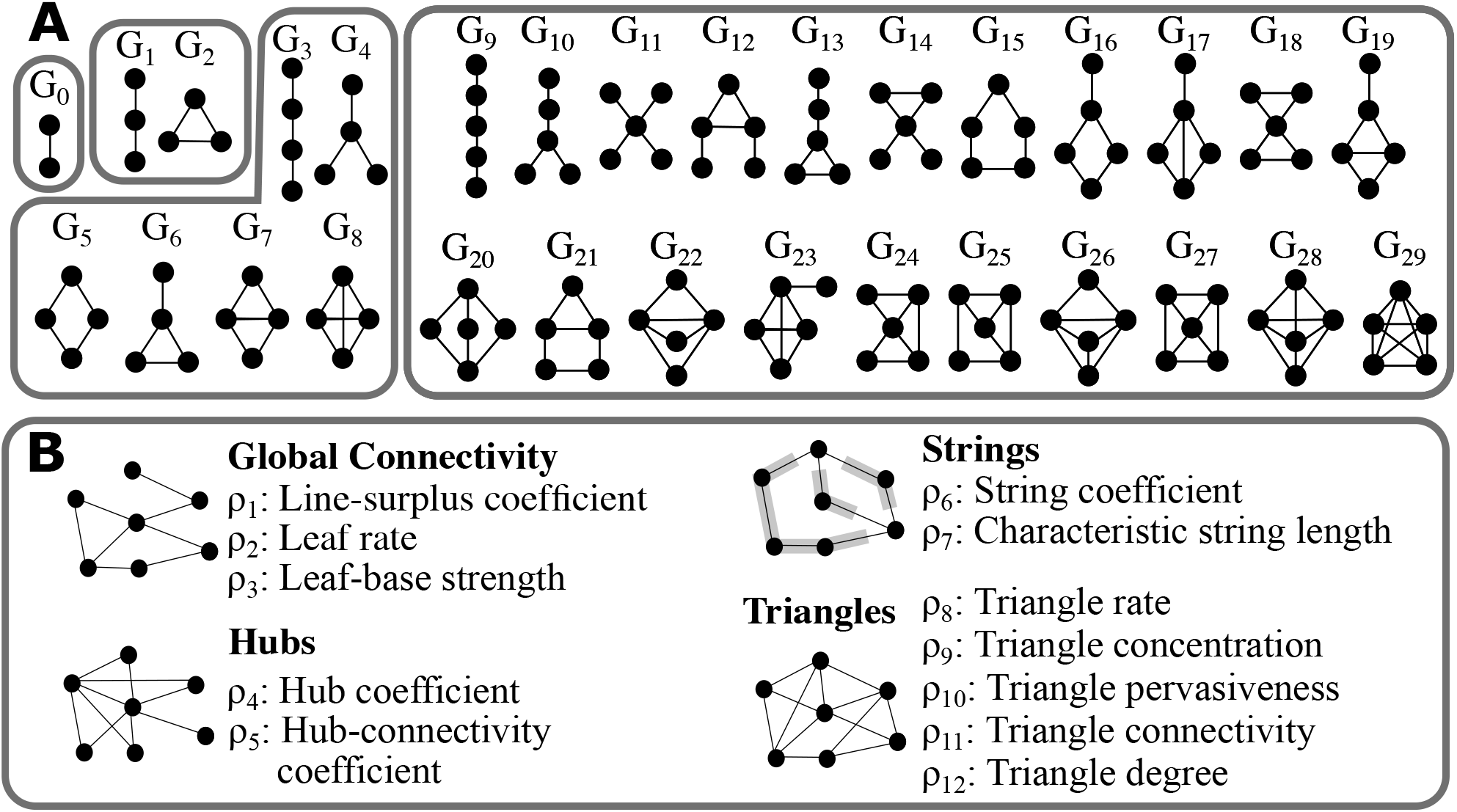
Vector representations of graph topology. (A) Undirected graphlets up to five nodes, organized by the number of nodes in the subnetwork. (B) GHuST coefficients.^8^ Refer to the original publication for formal definitions.

#### GHuST

While graphlets can be efficiently counted in a network, graphlet characterization can be biased when networks are different sizes and densities. Recent work by Espejo et al.^8^ developed the GHuST framework: twelve network statistics derived from two- and three-node graphlets (Fig. 1B). The GHuST framework captures both local and global network topology without needing four- and five-node graphlets.

The GHuST framework involves calculating twelve network statistics (also called *ρ* coefficients) based on the orbits from the first three graphlets (*G*_0_ – *G*_2_). These coefficients are grouped into four types of characteristics. Global connectivity coefficients measure the proportion of additional edges beyond those required for connectivity as well as leaf proportion and distribution. Hub coefficients measure the proportion and distribution of hubs in the network. String coefficients measure the number and proportion of consecutive nodes of degree two (consecutive *G*_1_ graphlets). Finally triangle coefficients measure the proportion, distribution, and connectivity of triangles (*G*_2_ graphlets) in the network. For simplicity, we call the *ρ* values *GHuST coefficients* and we have implemented them in our software.

## 3. Topological Structure of Pathway Databases

We asked firstly whether pathways are enriched for particular graphlets or GHuST coefficients within databases, and secondly whether these pathways are distinguishable from random sub-graphs of a protein-protein interactome. While previous work has compared graphlet distributions to some random models,^18,28^ we used two random graph models that are particularly well suited to answer our questions.

### 3.1. Graphlet Enrichment by Random Rewiring

To determine whether pathways are enriched for graphlets or GHuST coefficients, we used a random rewiring random graph model (REWIRE). In the REWIRE model, pairs of edges from a graph *G* are randomly rewired, preserving the degree sequence of *G*. For each pathway *p*, a REWIRE realization rewires on average ten times the number of edges in *p*; we generated 100 realizations of the rewire model. For each graphlet or GHuST coefficient, we computed the Z-score of the observed value compared to the values from 100 REWIRE realizations and counted the number of pathways with values that were larger than two standard deviations from the average. Specific graphlets and GHuST coefficients exhibit statistically significant over or under-representation in the Panther database (Fig. 2). The rewire null model networks have the same number of nodes and edges as the original networks, thus, the first graphlet (number of edges, *G*_0_) and the first GHuST coefficient (line-surplus coefficient *ρ*_1_) are not significant by constuction. However, many other structural properties that show discernible pattern in pathways are prevalent (Fig. 2), and these non-random features may be utilized to characterize pathways. Not all graphlet counts are independent of each other;^28^ for example, *G*_2_ can be calculated from *G*_0_ and *G*_1_, and this redundancy can be seen in the over-representation of *G*_4_ and *G*_1_1 (Fig. 2). GHuST coefficients are derived two- and three-node graphlets (e.g. the triangle rate *ρ*_8_ is closely related to the number of triangle *G*_2_), and many triangle-based *ρ* values are similarly over- or under-represented. Strikingly, pathway databases exhibit unique graphlet and GHuST enrichment patterns (Supplementary Fig. S2 and S3).

**Fig. 2.**
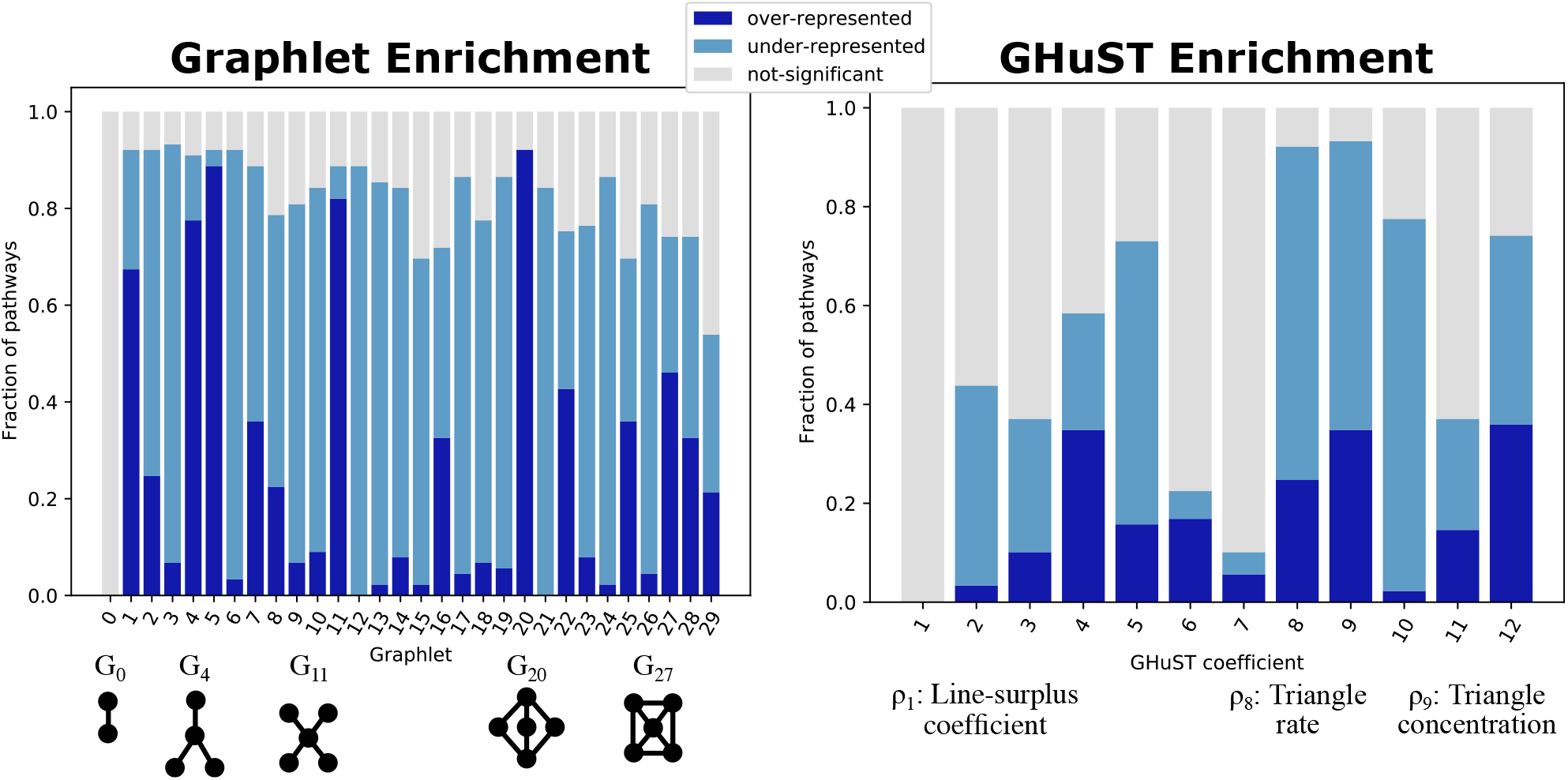
Over- and under-representation of graphlet counts (left) and GHuST coefficients (right) in the Panther Database. Select graphlets and *ρ* values are labeled below the x-axes.

### 3.2. Distinguishing Pathways from Random Subnetworks

Next, we considered whether the pathways were distinguishable from other subnetworks of a protein-protein interactome from PathwayCommons (Table 1). We wanted to preserve pathway connectivity and size when extracting a subnetwork from the interactome. To do so, we designed a random walker induced random graph model (walker). The walker model works as follows: given an interactome *G* and a pathway *p* containing *n* nodes and *m* edges, select a random node from *G* and perform a random walk until *n* nodes have been visited. Then, take the induced subgraph of *G* given the visited nodes to get subnetwork *H*. At this point if *H* has *m* or fewer edges, return *H*. If not, remove edges from *H* at random until *H* has *m* edges as long as their removal would not create a connected component of size 1. The walker-sampled networks are (in practice) the same size as *p*.^*^ To evaluate how well pathways from a database are distinguishable from the walker model we randomly generate one realization for every pathway in the database, thus building a balanced dataset with the same number of walker graphs as empirical pathways. We do this five times to generate five balanced datasets.

In this and the remaining analyses we cluster the vector representations of the pathways (either 30-dimensional graphlet vectors or 12-dimensional GHuST vectors). We perform agglomerative clustering with a mean linkage criterion and a cosine distance metric. Clustering quality is quantified using adjusted mutual information (AMI), which adjusts for random chance. Given a partition *X* determined by the agglomerative clustering and correct labels *Y* (here, “pathway” or “walker”), the AMI is defined as

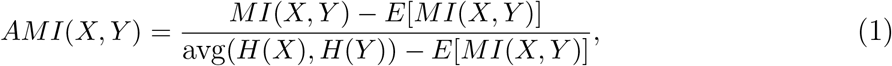

where *MI*(*X*, *Y*) is the mutual information of the partitions, *E*[*MI*(*X*, *Y*)] is the expectation of the mutual information of two partitions based on a hypergeometric model of randomness, and *H*(*X*) is the entropy of *X*. A larger AMI indicates that the partitions are more similar, and hence *X* better reflects the correct labels *Y*. We calculate the AMI for every possible number of clusters admitted by the agglomerative clustering algorithm.

When clustering the balanced datasets, pathways in each database are distinguishable
from walker networks, with larger AMIs associated with fewer clusters. The dendrogram of clusters by graphlet counts for the NetPath database, for example, contains only three NetPath pathways grouped with random networks in an otherwise perfect clustering (Fig. 3A). The AMI of this dendrogram reflects the good clustering, especially with few clusters since we have two labels (Fig. 3B left). To compare these results to other topological features such as clustering coefficient, we took a single dimension from each of the graphlet counts and GHuST coefficients that captured this information (*G*_2_ and *ρ*_8_, respectively) and clustered the same networks using Euclidean distance; the AMI for both metrics were notably worse (Fig. 3B middle). When clustering all 1,592 pathways and their walker models, we find that graphlets and GHuST coefficients cluster well in aggregate (Fig. 3B right). Supplementary Fig. S4 and S5 contain AMI plots for all individual databases.

**Fig. 3.**
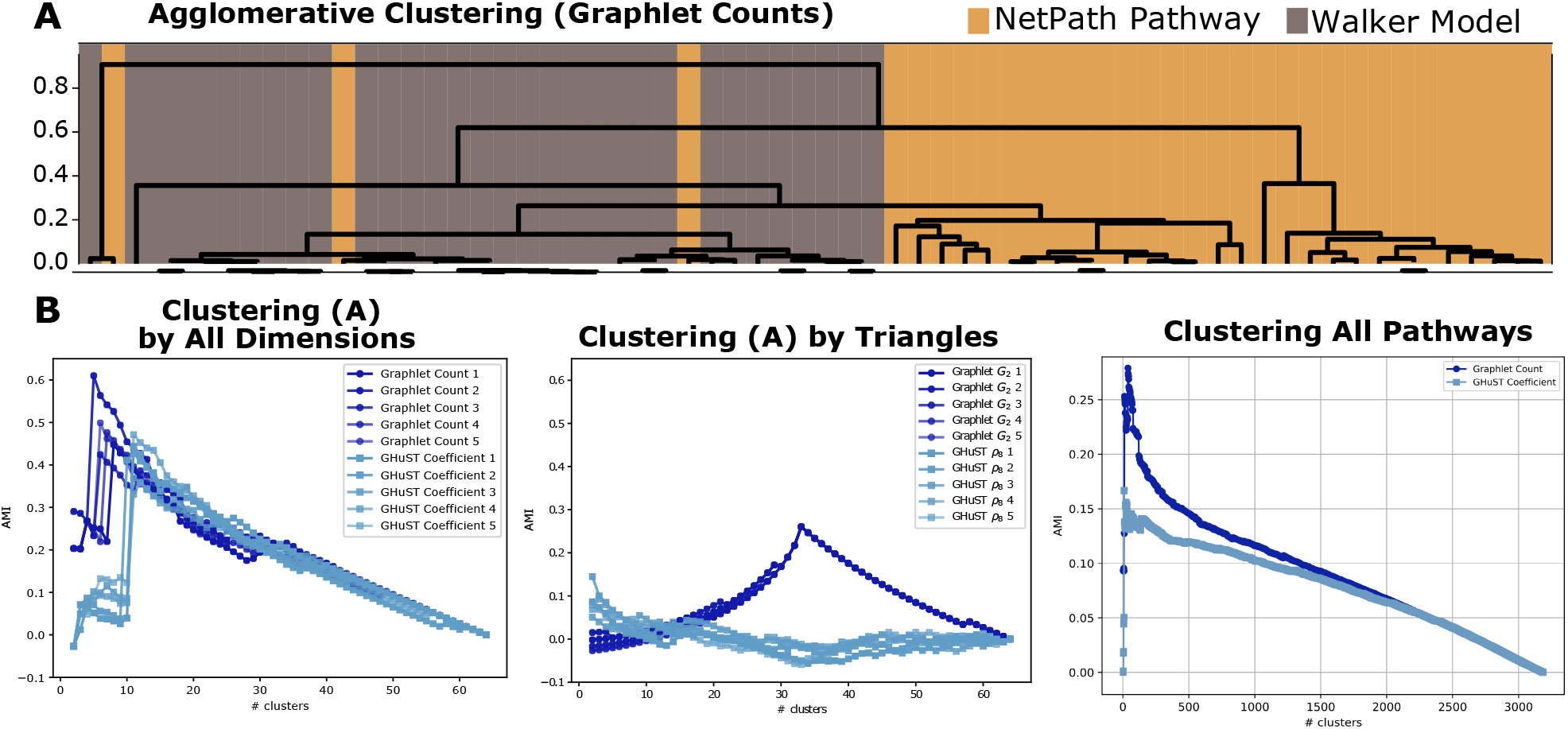
(A) Clustering of 32 Netpath pathways (orange leaves) and 32 WALKER networks (brown leaves) by graphlet counts. (B) AMI of five balanced datasets from NetPath using graphlet counts and GHuST coefficients. (C) AMI of five balanced datasets from NetPath using *G*_2_ and *ρ*_8_. (D) AMI of the balanced dataset containing all 1,592 pathways using graphlet counts and GHuST coefficients.

## 4. Topological Structure of Corresponding Pathways

After establishing that pathways are distinguishable from random graph models, we moved to directly comparing pathways across databases. To do so, we first need to identify a subset of pathways across the seven databases (INOH, KEGG, NetPath, Panther, PathBank, PID, and SIGNOR) that describe similar processes. We call these *corresponding pathways*, and we say that two pathways from different databases correspond if they aim to capture similar signaling events.

### 4.1. Identifying Corresponding Pathways

We employ a semi-automated procedure to find corresponding pathways from the seven databases. Our approach is similar in spirit to that of ComPath,^6^ which generates mappings between pathway annotations by considering the lexical similarity between names and content similarity between genes of each pair of pathways followed by a manual curation. Given two pathway databases *A* and *B*, pathway *a* ∈ *A* and *b* ∈ *B* are corresponding pathways if two conditions hold.

1. Tokenized versions of *a*’s and *b*’s pathway names share at least one word, after ignoring domain-specific terms (e.g., signaling, activation, network, downstream, etc.) and common stop words.
2. The asymmetric Jaccard overlap *J*(*a*, *b*) is non-zero; that is, *a* and *b* have at least one node in common (normalized by the number of nodes in *a*).

Let *f*(*a*, *B*) denote the set of pathways *b* ∈ *B* that correspond with pathway *a*. Two pathways *a* ∈ *A* and *b* ∈ *B* are *symmetrically corresponding* if they are corresponding pathways and each have the largest asymmetric Jaccard overlap among all other corresponding pathways in each database:

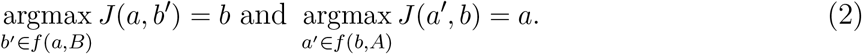

Since we are trying to find broad canonical pathways across databases we ignore pathways that include diseases or non-canonical terms (e.g., cancer, syndrome, viral, inflammation, etc.), model organisms (e.g., xenopus, drosophila, etc.), or metabolic signaling terms.

Next, we must identify groups of symmetrically corresponding pathways that collectively describe a single event across multiple databases. To do so, we build an undirected graph *G* = (*V*, *E*) where the nodes are pathways and two nodes are connected if the pathways are symmetrically corresponding (Fig. 4A). We find connected components in *G* that contain pathways from at least *τ* different databases, where *τ* is a user-defined threshold (we use *τ* = 6).

**Fig. 4.**
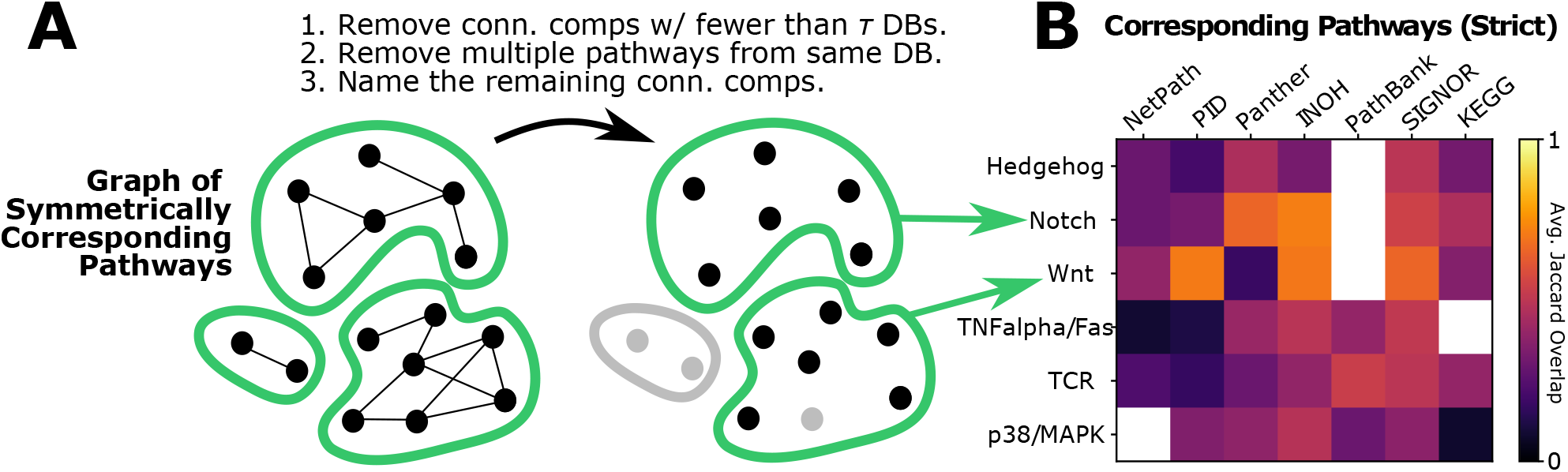
(A) Identifying corresponding pathways. We first build a graph where the nodes are pathways and two edges denote symmetric correspondence. Then, we find the connected components that contain at least *τ* different databases and ensure at most one pathway per database. Finally, we name each connected component based on the pathway names. (B) Each connected component is represented as a row in the matrix, which describes the average Jaccard overlap of each pathway across databases. White entries denote databases with no corresponding pathway.

Finally, we have a last manual step that examines each connected component that passes the *τ* threshold, assigns a common name to the pathway, and selects exactly one pathway for each database based on the pathway name.^†^ If we cannot determine a common name, we remove that connected component from consideration. Once we have a table of corresponding pathways for each database, we add the KEGG-collapsed and SIGNOR-collapsed datasets, since they will have an exact match with the KEGG-expanded and SIGNOR-expanded titles. Complete details about gathering, parsing, and finding corresponding pathways are provided in the GitHub repository.

#### Corresponding pathways have low node overlap

While we used Jaccard overlap to determine corresponding pathways, this overlap was typically quite low. For each pathway, we calculated the Jaccard overlap for pairs of databases, resulting in a database-by-database heatmap of overlaps (Supplementary Fig. S6). We then calculated the average Jaccard value for each pathway/database combination (Fig. 4B). For Hedgehog, TNFalpha/Fas, TCR, and p38/MAPK, on average about a third of the nodes were shared between any two pathways. The Notch and Wnt pathways had slightly higher overlap, with an average of 0.5 across the rows. Notably, many of the overlaps were bleak, with minimums ranging from 0.08 (p38/MAPK) to 0.29 (Notch) across the rows.

#### Corresponding pathways cluster by database

Once we had corresponding pathways, we calculated the AMI from the agglomerative clustering based on cosine similarity as described in Section 3.2. We found that the AMI was much higher when we labeled partitions by database instead of by pathway (the blue “Original” curves in Fig. 5A). This indicates that there are database-specific topologies that are driving the clustering; we call this *database-specific structure*.

**Fig. 5.**
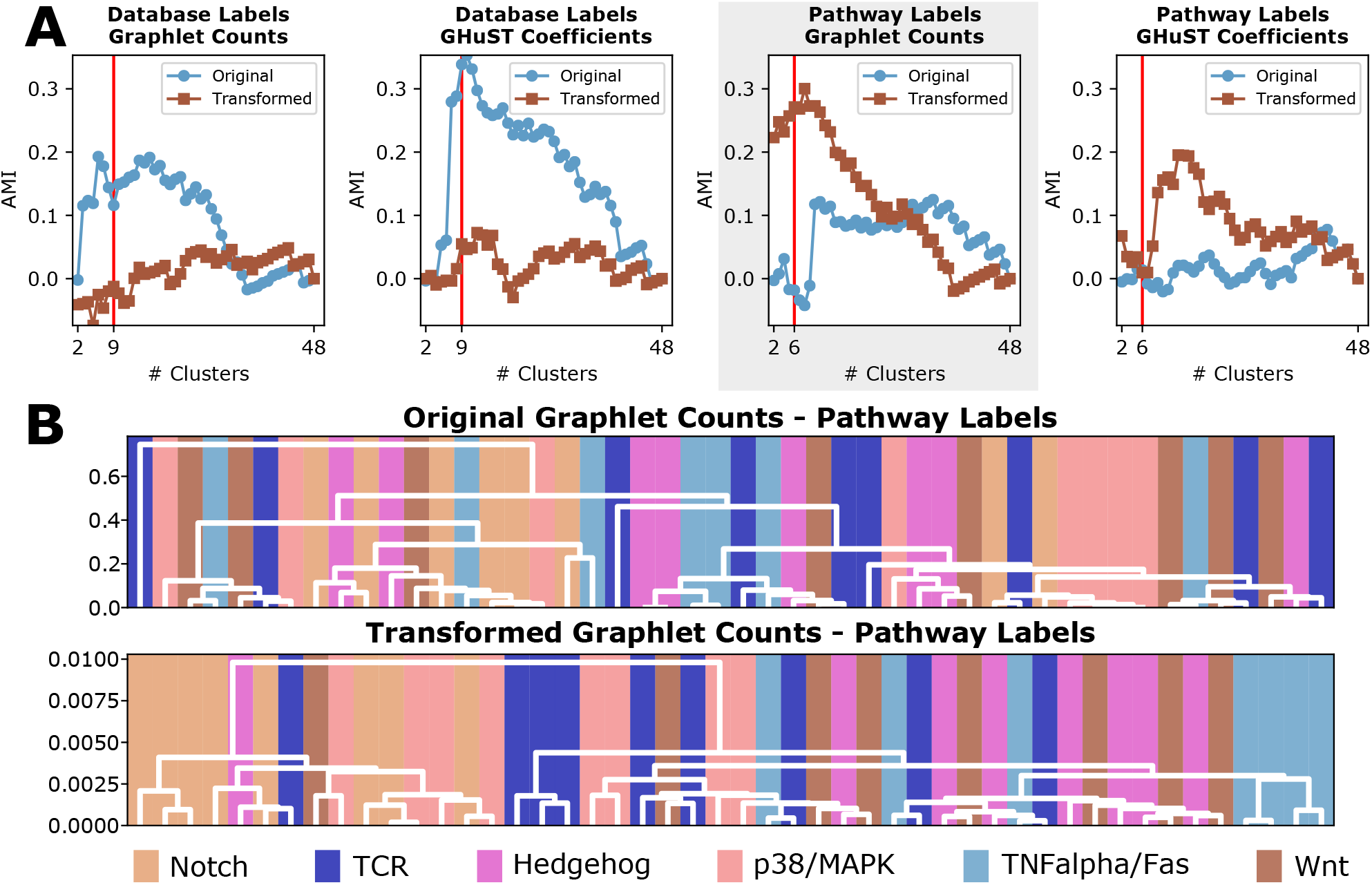
(A) AMI of graphlet counts and GHuST coefficients when clustering corresponding pathways using databases as ground truth labels (first two plots) or pathways as ground truth labels (last two plots). Blue lines indicate clustering by original values; brown lines indicate clustering by regression-transformed coordinates. Vertical red line indicates the correct number of clusters for each ground truth dataset. (B) Dendrograms of the clusters using original (top) and transformed (bottom) graphlet counts, colored by the six pathway labels. The AMI of these dendrograms are shown in the shaded plot in Panel (A).

## 5. Correcting for Database-Specific Structure Reveals Pathway Similarities

The blue “Original” curves in Fig. 5A suggest that pathways within a specific database share some topological similarities. This is not particularly surprising, since each database is designed by curators with different goals in mind. We wondered whether we could correct for the database-specific structure to reveal pathway specific topological features. To do this, we used a simple ordinary least squares (OLS) model to find database weights to transform pathway vector embeddings. For this part, we normalize the graphlet counts by all counts for graphlets with the same number of nodes (e.g., the number of triangles is normalized by the sum of *G*_1_ and *G*_2_). We treat each graphlet value or GHuST coefficient separately. Let 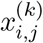 be the *i*th co-ordinate in the vector embedding where *j* denotes the pathway and *k* is the database label, in a set of at least six corresponding pathways. For each coordinate *i* and pathway *j*, we construct a profile *y* as an average over all databases as

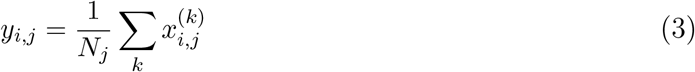

where *N_j_* is the number of databases that contain pathway *j*. Note that the average profile *y* is not specific to a database and only has two indices *i* and *j*.

We use the following linear regression for each coordinate *i* to identify database-specific structure within the pathway profiles *x* using *y* as the target function.

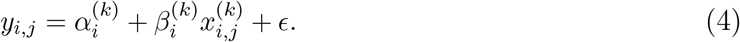

The estimated values of the intercept *α* and the regression coefficients *β* that minimize the error term *∈* can be used to transform any pathway profile in a database. The transformed profiles 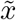 are computed as

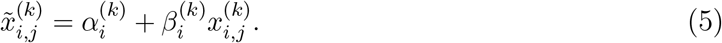

Recall that our goal is to have corresponding pathways cluster together, rather than databases. Clustering the transformed coordinates 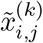 dramatically reduces the AMI for database labels while increasing the AMI for pathway labels (brown “Transformed” curves in Fig. 5A). This is illustrated with cluster dendrograms of the original and transformed graphlet counts (Fig. 5B). Not only does this illustrate that the database-specific structure is reduced, but pathways from different databases are closer in the transformed vector space. The effect of transformation can also be seen in the first two principal components of the GHuST coefficients, where some databases cluster in the original vector space and nearly all pathways cluster in the transformed vector space (Fig. 6 and Supplementary Fig. S9). Notch and Wnt overlap in the transformed space, which makes sense due to their extensive pathway crosstalk. The first two principal components of the transformed graphlet counts also reveals clustering by pathway (Supplementary Fig. S10).

**Fig. 6.**
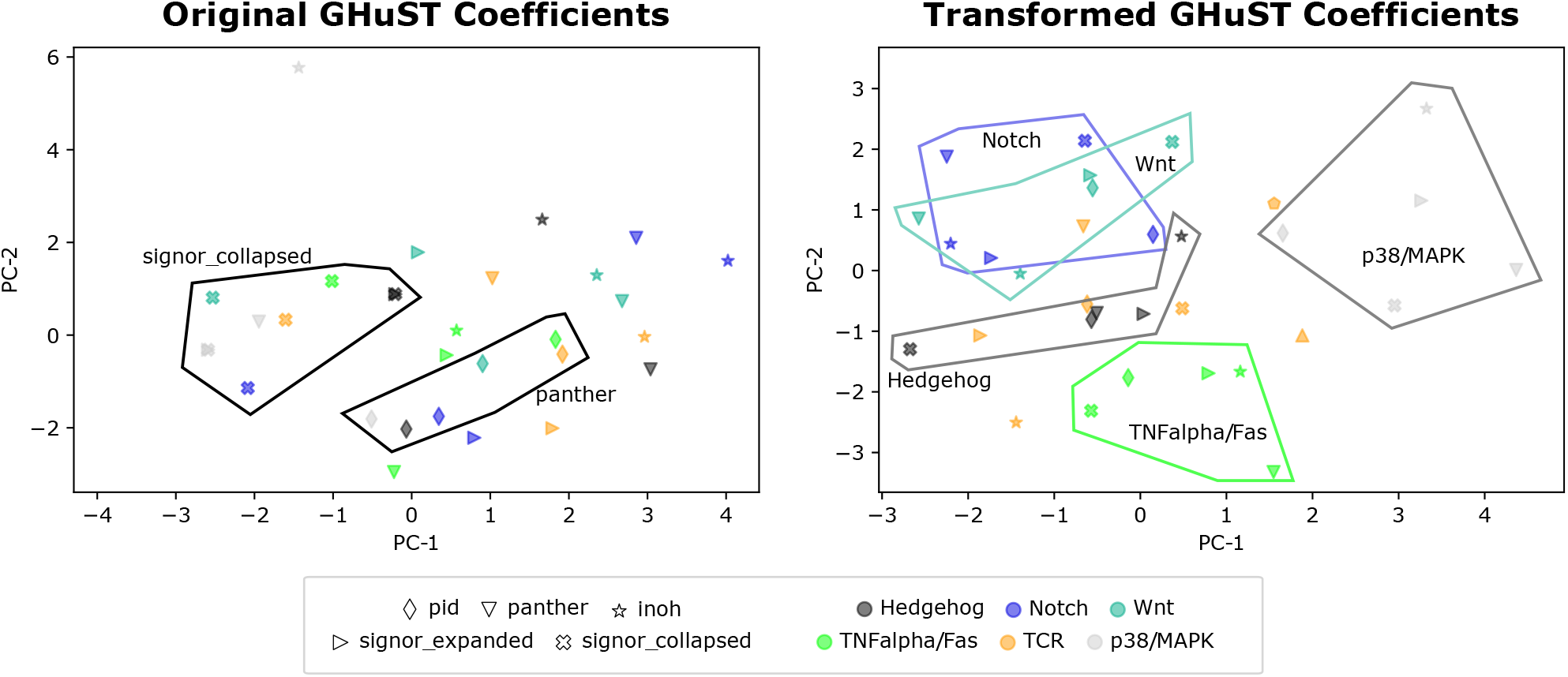
Principal component analysis (PCA) of the first two components for the original GHuST coefficients (left) and the transformed coefficients (right) for the five datasets that have all six corresponding pathways for *τ* = 6. Datasets are denoted by marker shape, pathways are denoted by colors. Representative clusters are annotated

## 6. Discussion

As the number of pathway databases grows, we have the opportunity to leverage more curated interactions to better understand cellular behavior and response. However, combining signaling pathways across databases is not a trivial task, and databases may each have structural features that obfuscate latent pathway structure. We have presented a topology-based framework for describing pathway structure and reconciling signaling pathways across databases. Signaling pathway structure is consistently distinct from random graph models, even when accounting for node degree and comparing to appropriate subnetworks from a larger interactome. Using a new approach that accounts for database-specific structure, we show that corresponding pathways cluster together and ultimately share topological features despite coming from differently-curated resources.

We use two embeddings based on graphlets to describe network topology. We found the AMI curves based on graphlet counts were often slightly larger than those based on GHuST coefficients, but it is striking that GHuST performs so well given that the coefficients are derived from only two- and three-node graphlets. Further, neither embedding worked particularly well when reducing *τ* to four for corresponding pathways, though the trends of the original and transformed AMIs still hold (Supplementary Fig. S7, S8, and S11). We suspect there is room for more descriptive statistics using four node graphlets in a GHuST-like framework which may outperform both GHuST coefficients as well as graphlets for the study of signaling structure.

We (and the folks we build our work upon) have to make many choices in the design and execution of our work, and thus there are many sources of bias in our study. First and foremost, the foundation of our work builds upon the manual curation of different databases. Some databases are considerably more well-developed than others - KEGG, for example, is over twenty years old and is still actively maintained. Newer databases like SIGNOR and PathBank are smaller than others but are quickly growing. Researchers might be more familiar with a particular pathway database and slower to adopt new resources, which might also lead to bias in the peer review of database publications. For example, updates of existing databases may be more well-received than new databases that offer complementary resources.

Certain pathways are more studied than others, and canonical versions of pathways are more often described than non-canonical counterparts. Examples of well-studied pathways appear in our corresponding pathway lists, since we require that the pathways are present in multiple databases. This might expressly contradict the goal of particular pathway databases, for example PathBank includes model organisms beyond human pathways.

The decisions made by us and others to interpret biochemical reactions as graphs undoubtedly affects our results. Firstly, nearly all pathway databases capture directed, signed interactions, and standardized pathway formats like BioPAX and SBML capture multi-way relationships and reaction stochiometry. These details are ignored when we use undirected graph representations. We partially addressed this issue with the “expanded” versions of KEGG and SIGNOR pathways, but it certainly deserves more investigation. Directed graphlets^1,21^ and signed graphlets^4^ could reveal more refined pathway structure.

Our methods are intentionally straightforward, with the goal to show that using topological metrics on undirected networks can reveal pathway-specific structure. In addition to improving the underlying graph representation of signaling, there is room for improvement in the choice of clustering and the regression model for correcting database-specific topologies. Additionally, we note that the transformed coordinates do not directly translate into networks that exhibit those coordinates. An exciting area of future work is to identify subgraphs from a larger interactome that approximates an arbitrary graphlet count vector.

As the number of signaling pathway databases grows, topological features hold promise in elucidating pathway specific structure. While signaling pathway representations have long been acknowledged to be different within and across databases, we have shown that it is possible to reduce database-specific structure and find structural similarities among corresponding pathways. Our work indicates that reconciling pathways while retaining the databases as separate entities can characterize signaling pathway structure.

## Data Availability

https://github.com/Reed-CompBio/pathway-reconciliation.

## Code Availability

Code to parse databases, count graphlet and calculate GHuST coefficients, and generate all results is available at https://github.com/Reed-CompBio/pathway-reconciliation.

## Acknowledgments

This work is funded by the National Science Foundation (DBI #1750981) to AR.

## Footnotes

* This works because the wALKER-induced subgraphs tend to be much denser than pathways, and it is generally straightforward to find edges which are removable without creating isolated nodes.

† Note that it is possible that a connected component may have two pathways *b*, *b′* ∈ *B* from the same database if they were symmetrically corresponding with other pathways in different databases.

