## Supplementary Information for "Stop Bickering! Reconciling Signaling Pathway Databases with Network Topologies"

***Supplementary Material for***
### Stop Bickering! Reconciling Signaling Pathway Databases with Network Topologies

Tobias Rubel*, Pramesh Singh*, and Anna Ritz†

*Biology Department, Reed College, Portland, Oregon, USA*

*Equal author contribution

†

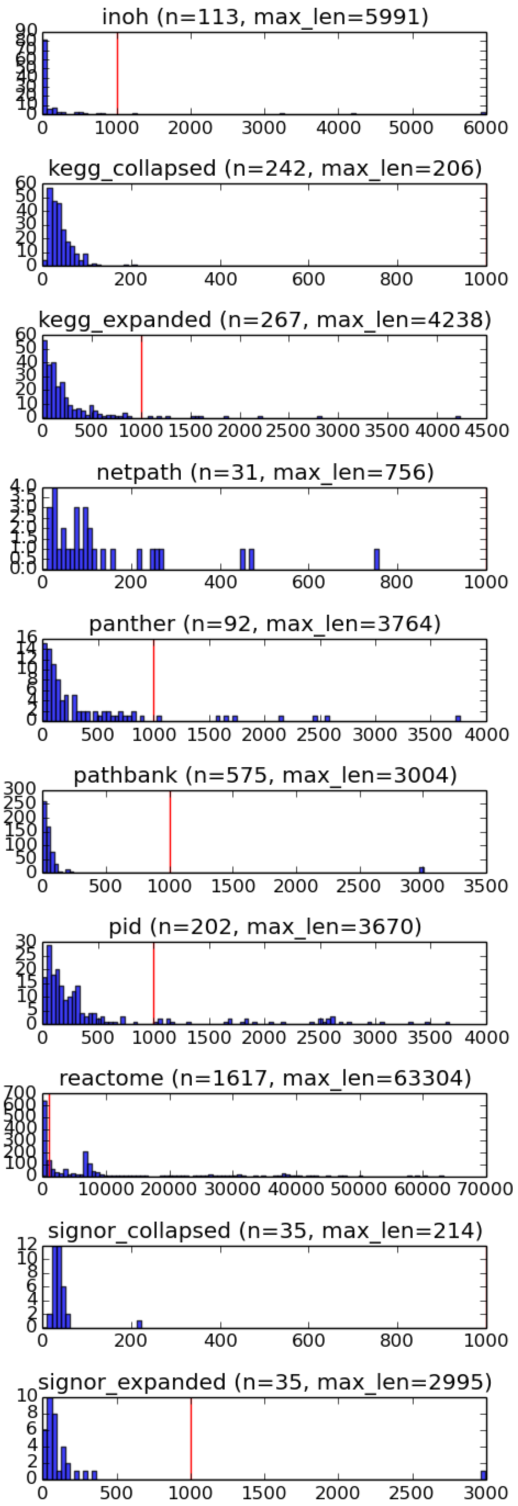


Fig. S1. Histograms of pathway sizes (by number of interactions) for all datasets considered. Vertical red bar indicates clusters with 1,000 interactions. Reactome pathways are about two orders of magnitude larger than other pathway databases. Additionally, the “-expanded” pathways are larger than the “-collapsed“ pathways, as expected.

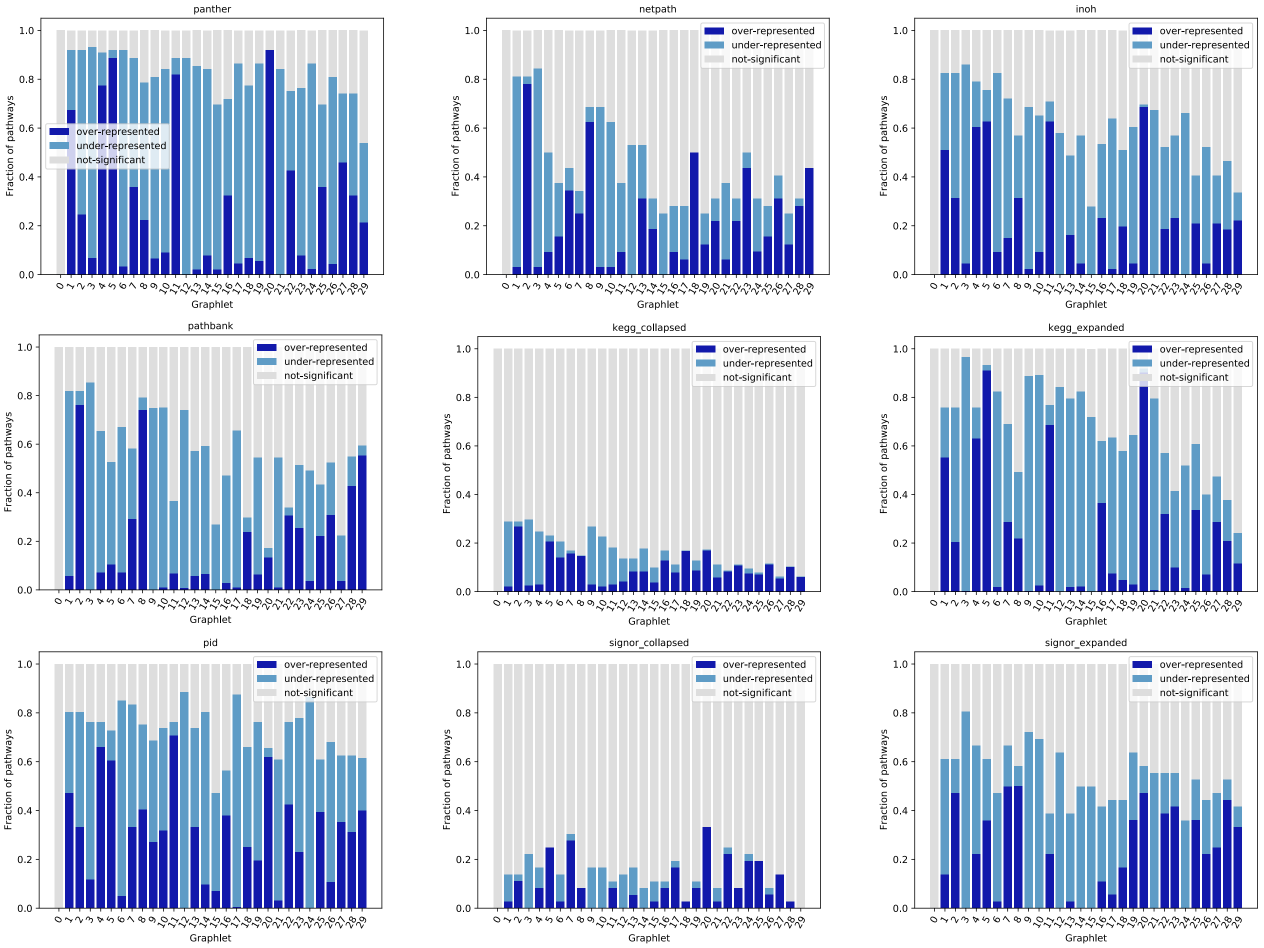


Fig. S2. Over- and under-represented dimensions of the 30 graphlet counts for all datasets. The Panther pathway database is shown in the main paper. Note that the "-collapsed" versions of KEGG and SIGNOR, which include protein complexes and families as nodes, have many fewer over- or under-represented dimensions.

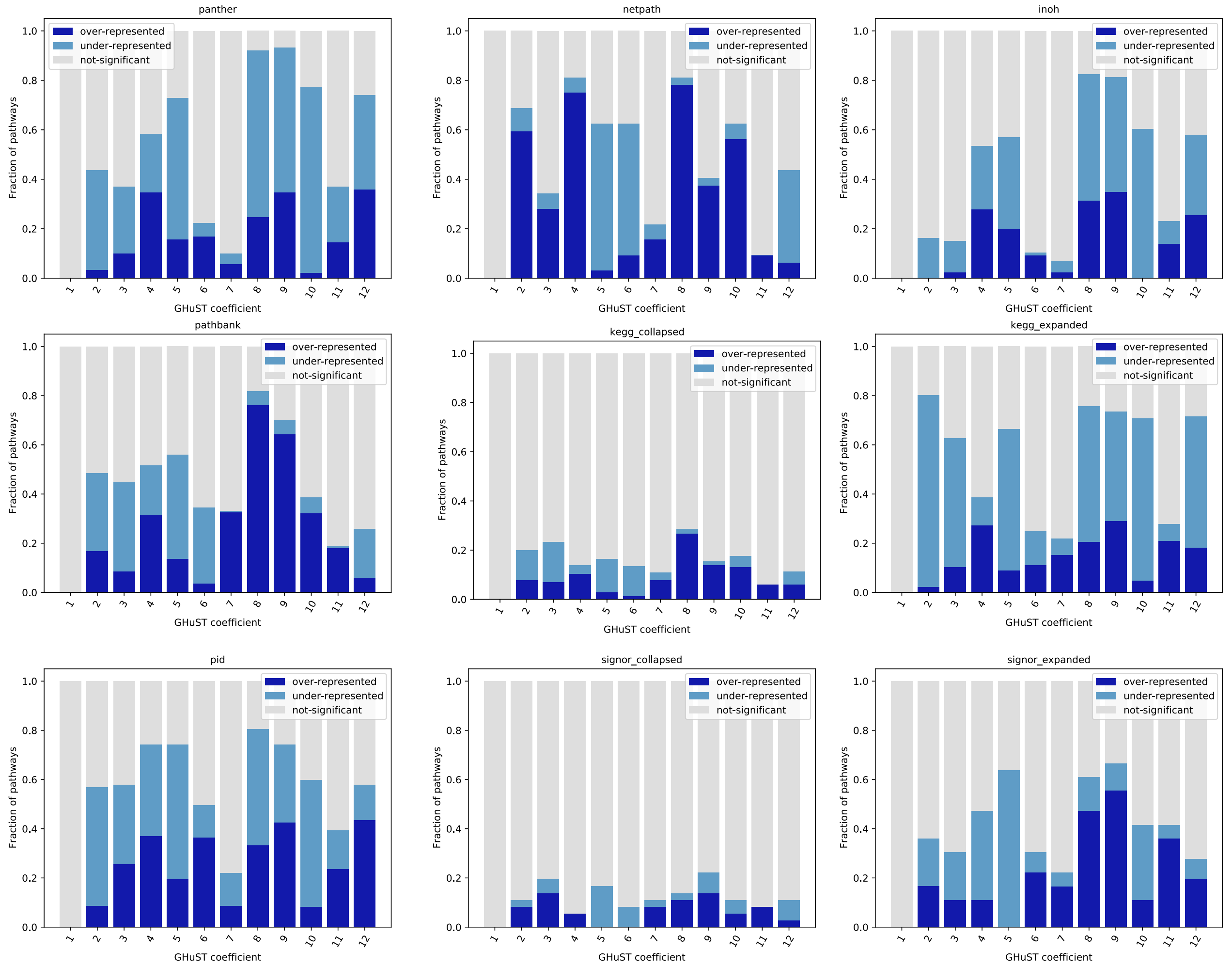


Fig. S3. Over- and under-represented dimensions of the 30 GHuST ($\rho$) coefficients for all datasets. The Panther pathway database is shown in the main paper. Note that the "-collapsed" versions of KEGG and SIGNOR, which include protein complexes and families as nodes, have many fewer over- or under-represented dimensions.

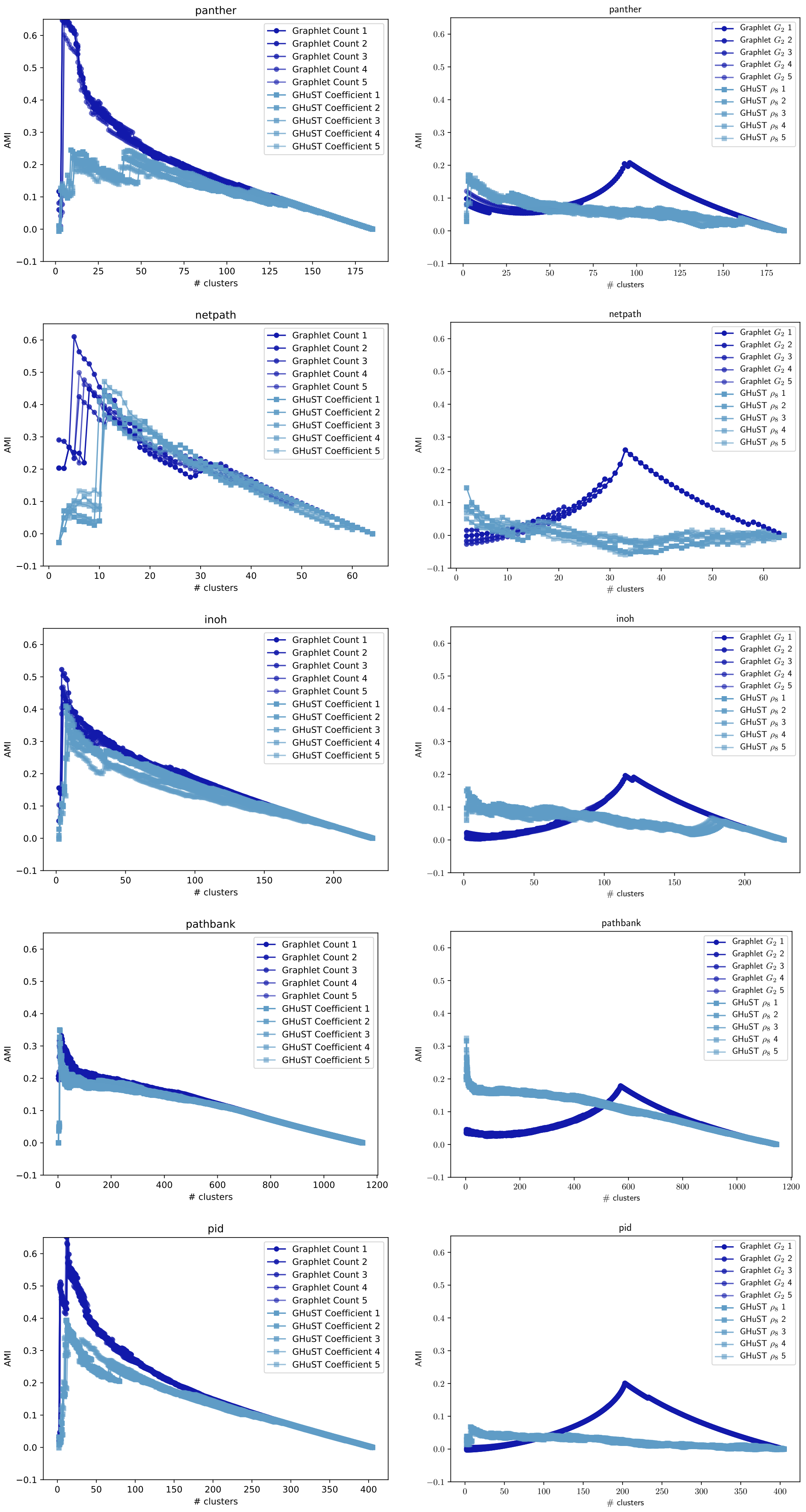


Fig. S4. AMI plots for full vectors (left) and triangles only (right) for pathway databases.

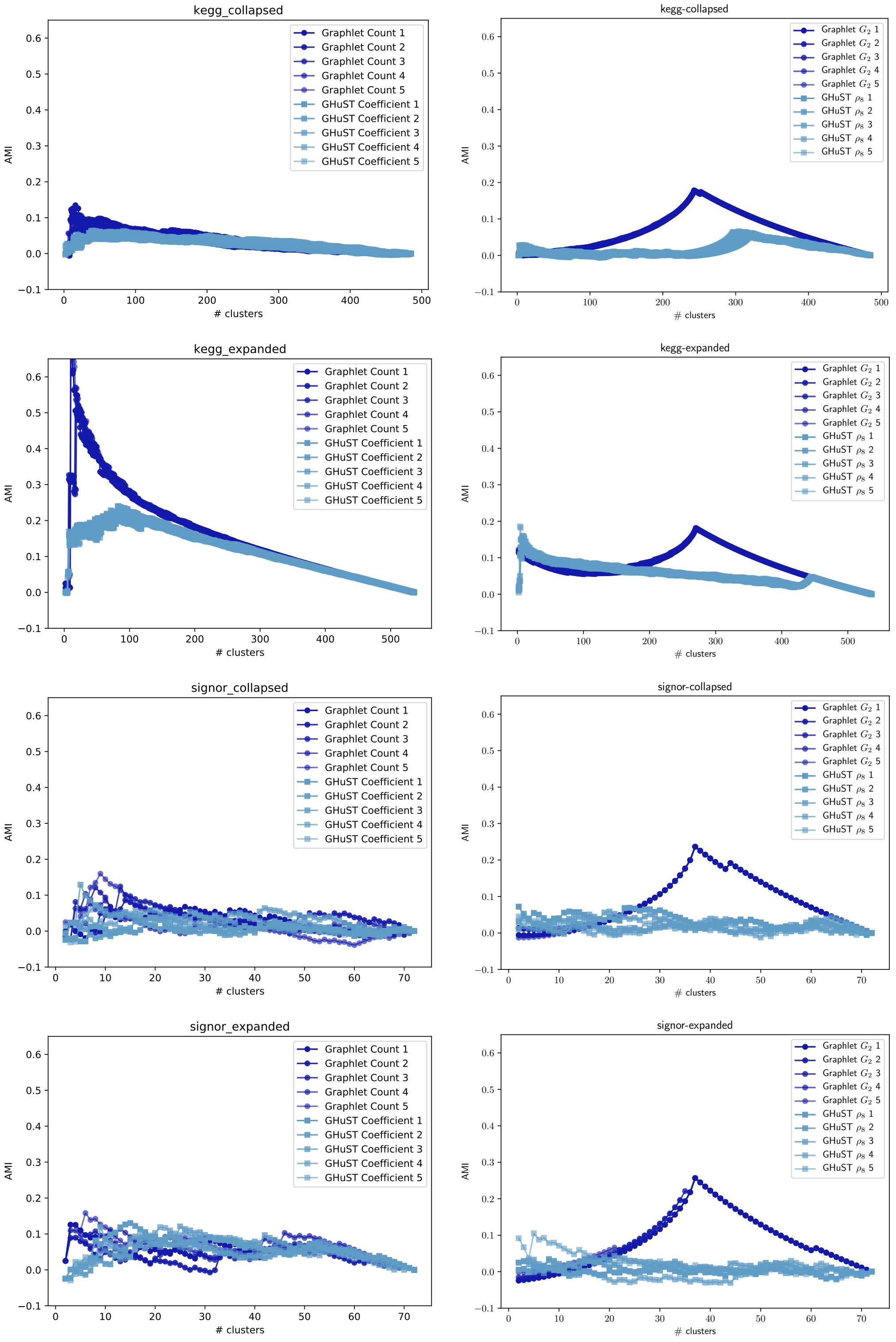


Fig. S5. AMI plots for full vectors (left) and triangles only (right) for pathway databases.

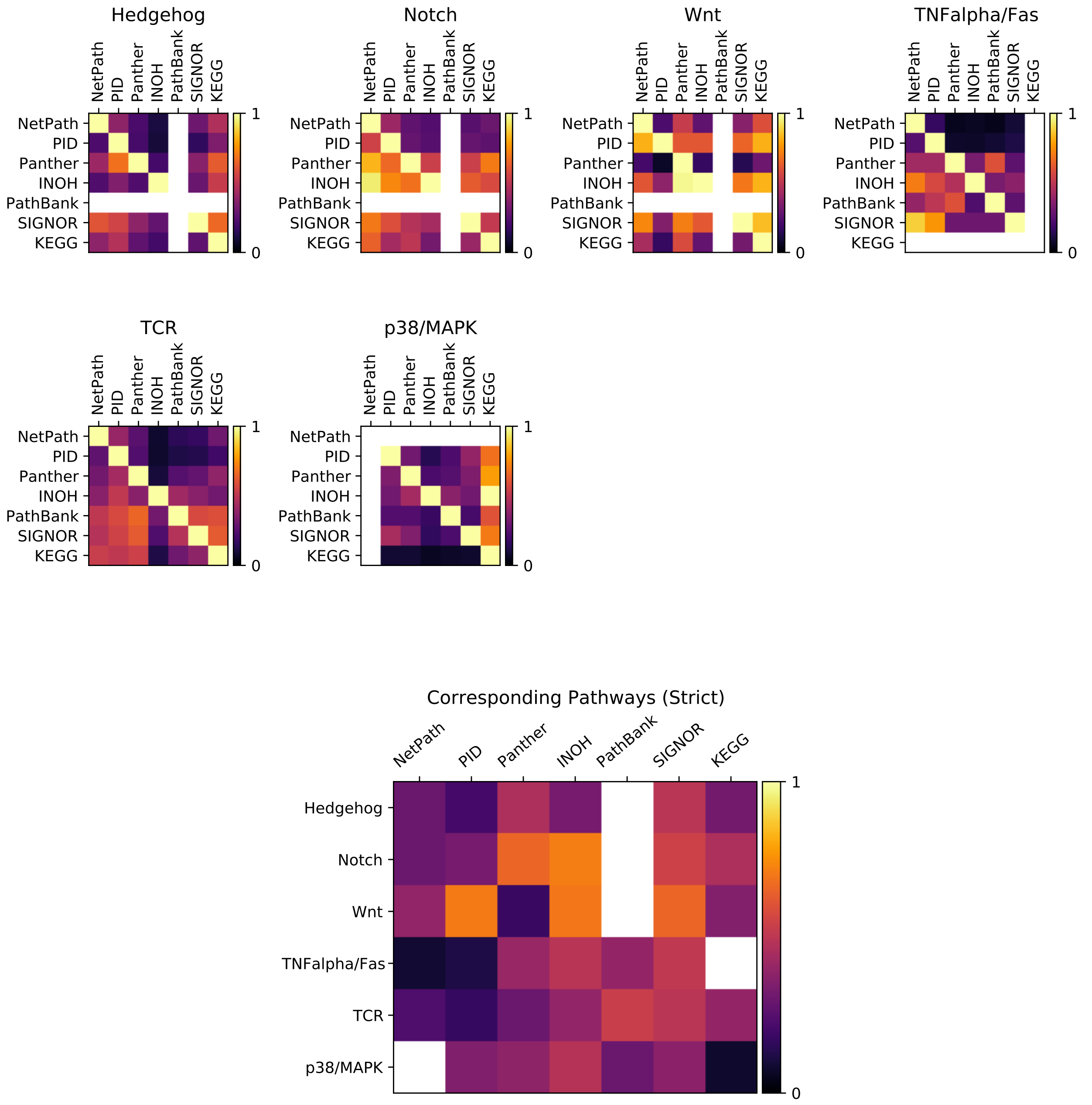


Fig. S6. Asymmetric Jaccard overlap of nodes across databases for each pathway (top) and averaged (bottom) when $\tau = 6$. White entries denote databases that do not have corresponding pathways. In the bottom figure, all non-identity and non-missing entries of the rows are averaged.

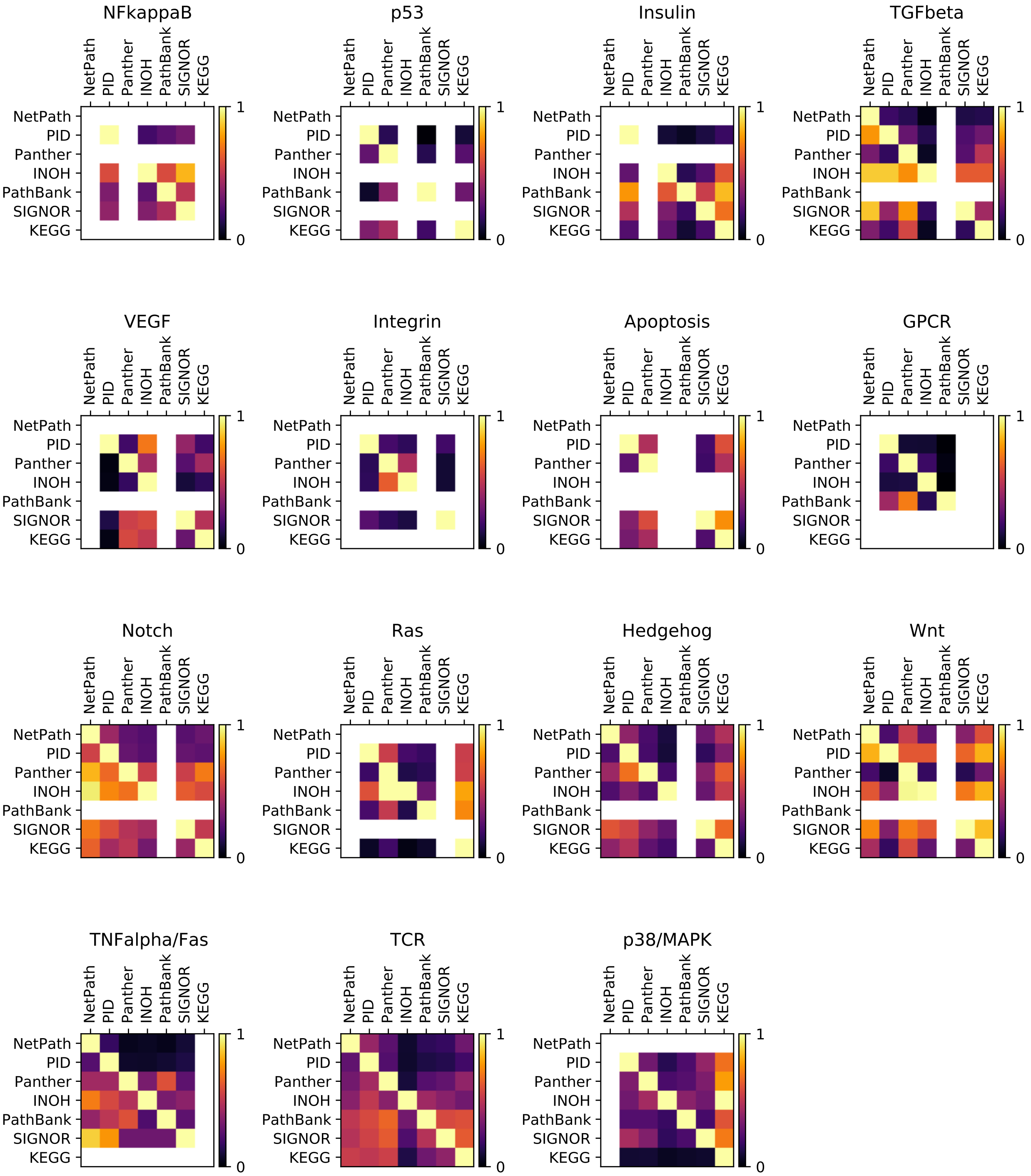


Fig. S7. Asymmetric Jaccard overlap of nodes across databases for each pathway when $\tau = 4$. White entries denote databases that do not have corresponding pathways.

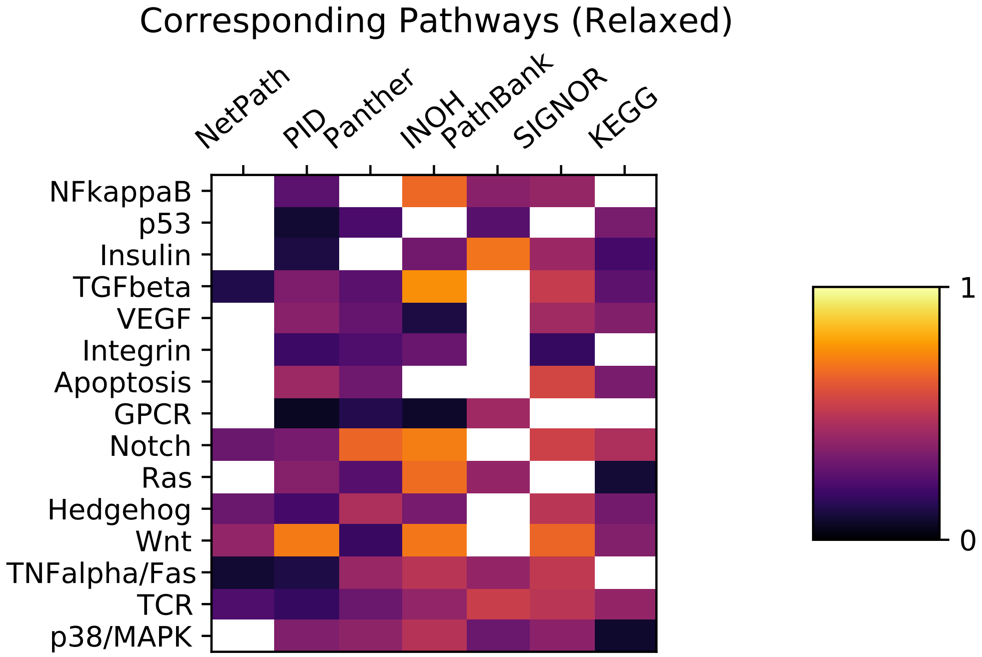


Fig. S8. Asymmetric Jaccard overlap of nodes averaged across databases for each pathway in the relaxed scenario (entries from Fig. S7). White entries denote databases that do not have corresponding pathways. All non-identity and non-missing entries of the rows are averaged.

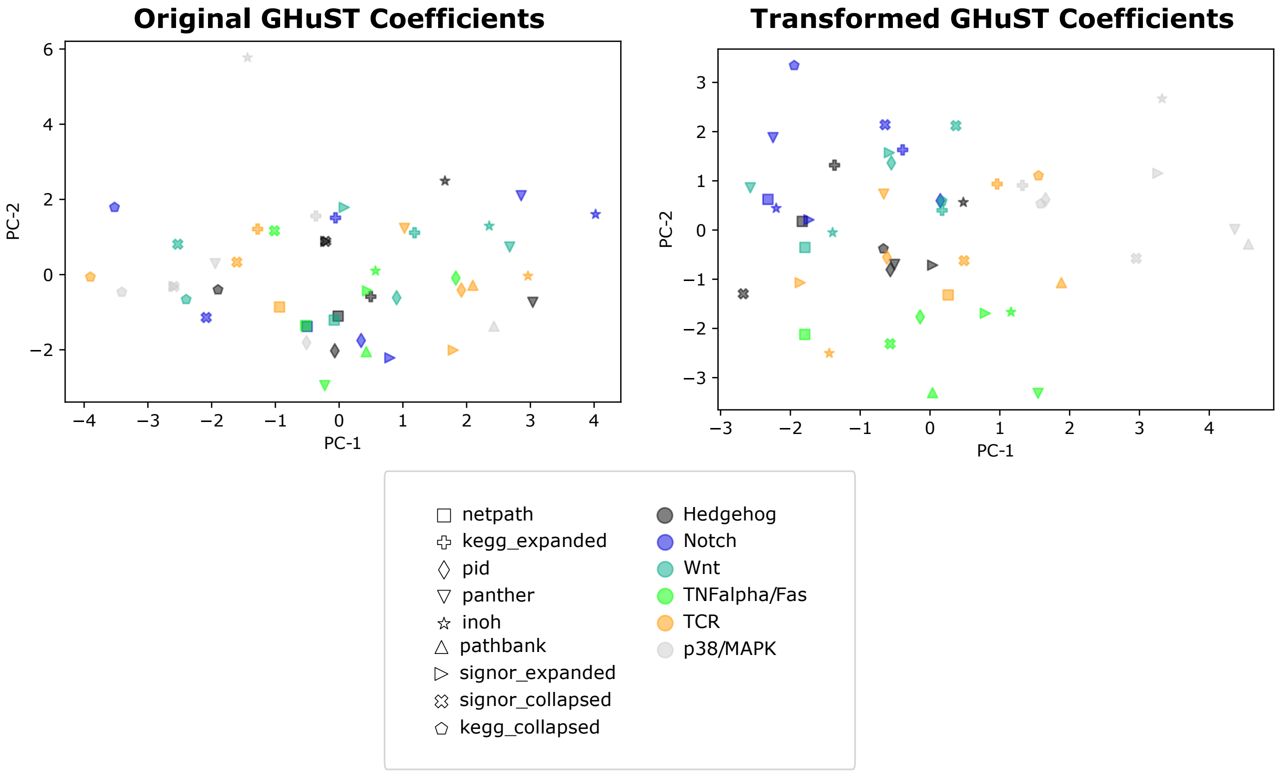


Fig. S9. Principal component analysis (PCA) of the first two components for the original GHuST coefficients (left) and the transformed coefficients (right) for all nine datasets. Datasets are denoted by marker shape, pathways are denoted by colors.

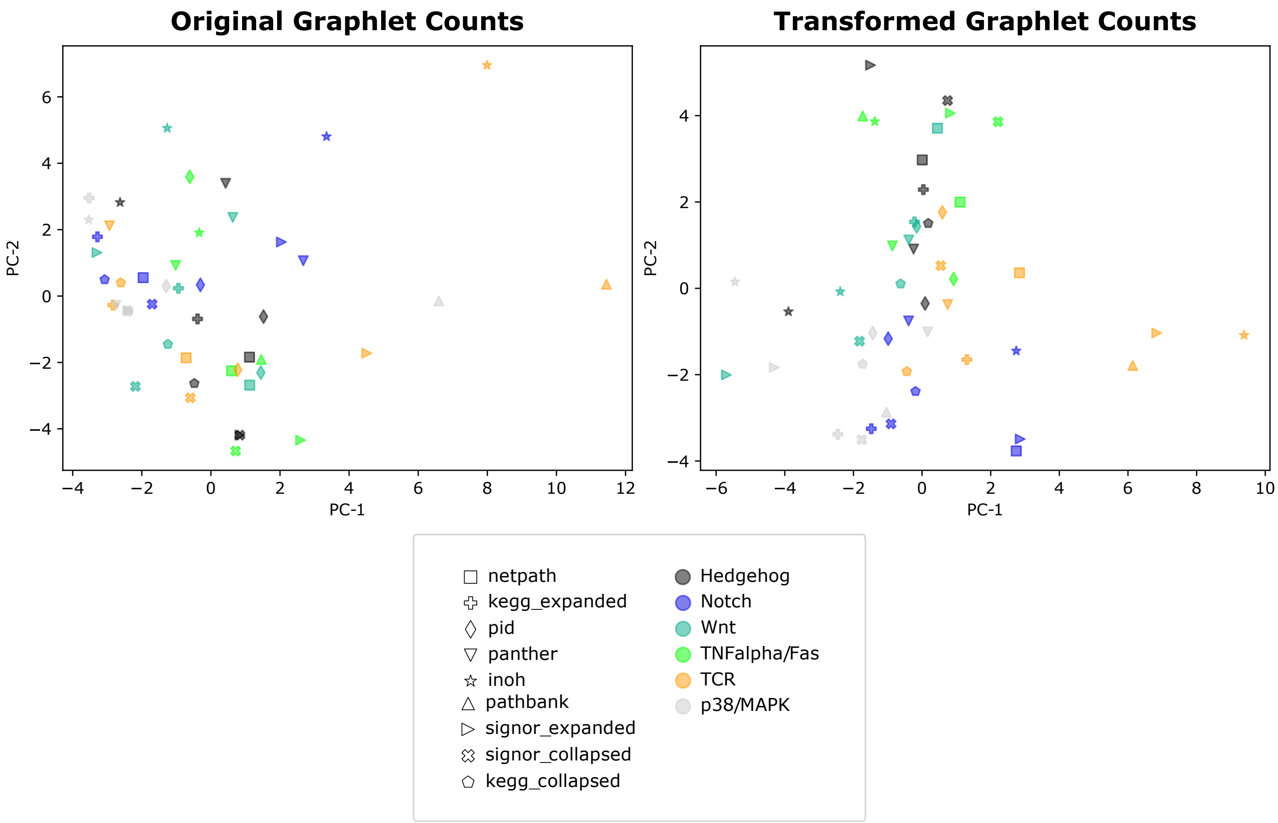


Fig. S10. Principal component analysis (PCA) of the first two components for the original graphlet counts (left) and the transformed coefficients (right) for all nine datasets. Datasets are denoted by marker shape, pathways are denoted by colors.

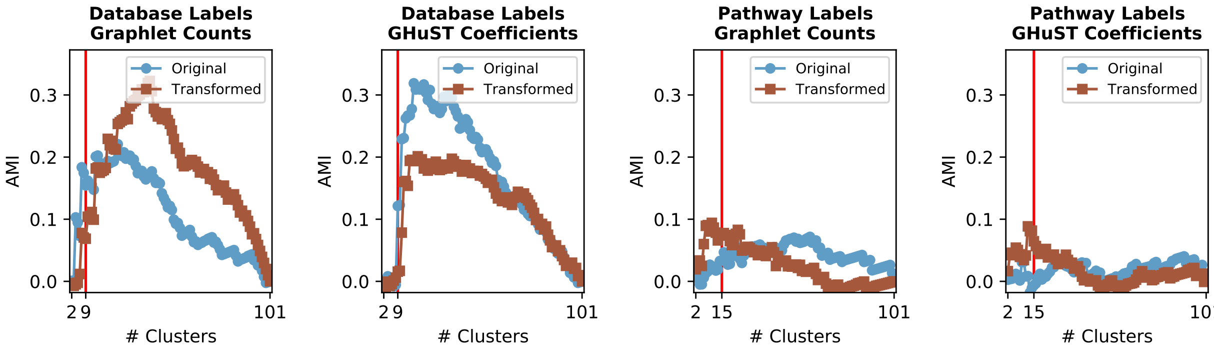


Fig. S11. AMI of graphlet counts and GHuST coefficients when clustering corresponding pathways ($\tau = 4$) using databases as ground truth labels (first two plots) or pathways as ground truth labels (last two plots). Blue lines indicate clustering by original values; brown lines indicate clustering by regression-transformed coordinates. Vertical red line indicates the correct number of clusters for each ground truth dataset.
